# Connectivity of Implicit Statistical Learning Networks is Associated with Language Impairment Severity in Post-Stroke Aphasia

**DOI:** 10.64898/2026.11.02.763658

**Authors:** Amy E. Ramage, Beau Underwood, Katelyn Lerner, Kirrie J. Ballard

**Affiliations:** University of New Hampshire, Department of Communication Sciences & Disorders, Durham, NH; University of New Hampshire, Interdisciplinary Program in Neuroscience & Behavior, Durham, NH; Faculty of Medicine and Health, The University of Sydney, Camperdown, NSW, Australia; The Brain and Mind Centre, The University of Sydney, Camperdown, NSW, Australia

**Author notes:** **Corresponding Author:** Amy E. Ramage, University of New Hampshire, Department of Communication Sciences and Disorders, 4 Library Way, Durham, NH 03824.

**Keywords:** implicit statistical learning, language, post-stroke aphasia, resting-state, functional connectivity

## Abstract

Rehabilitation of language for individuals with post-stroke aphasia (PSA) requires relearning to use cognitive-linguistic skills, but little is known about how these individuals relearn, or how brain reorganization may support different types of learning. One type of learning critical to language acquisition is implicit-statistical learning (ISL), which supports robust, accurate performance with minimal conscious effort. ISL relies on brain subnetworks that include language areas as well as domain general regions (i.e., salience and cognitive control) with the latter considered less likely to be damaged in individuals with PSA. Using three brain subnetworks previously found to be associated with ISL, the present study used graph theory metrics to (1) compare their functional connectivity in PSA (n = 30) relative to neurologically healthy controls (n=18), and (2) examine their relationship with aphasia severity in the PSA group. Resting-state functional connectivity was weaker for the language and salience subnetworks in PSA than controls, but not for the cognitive control subnetwork. Additionally, edges within all three networks correlated with aphasia severity, particularly between regions of the right frontal cortex, such that stronger connectivity in these ISL subnetworks were associated with milder aphasia. While it is unsurprising that the damaged left-sided brain regions in the PSA group resulted in connectivity that differed from controls, these differences were also observed in the salience subnetwork. Further, the correspondence to aphasia severity was evident in all subnetworks, highlighting that domain general ISL networks may contribute to variability in language outcomes.

## Introduction

Relearning the use of language is an important part of recovery in individuals with post-stroke aphasia (PSA). A long tradition of study that connects language performance to neural correlates has utilized the structure-function hypothesis (Geschwind, 1970), correlating lesion volume to symptom metrics (c.f., Levy et al., 2023). Growing from that approach, the connections between regions found to be associated with specific language functions have furthered the understanding of the complex processes of language. For example, the connectivity of brain regions involved in language, such as Broca’s and Wernicke’s areas, is known to be important for auditory perception and comprehension of spoken language (Hickok & Poeppel, 2007). However, investigation into the connectivity of each of these language areas separately elucidates a complex set of connections with a diffuse set of other cortical and subcortical structures (Tomasi & Volkow, 2012). This work has clarified the necessity of left hemisphere cortical structures for language processing but also indicates that they are not sufficient without contribution of the right hemisphere and subcortical structures (Copland et al., 2021; Hope et al., 2017; Ramage et al., 2020). The intricacies of the language network continue to be unveiled, but in aphasia recovery, the location of the lesion in the left hemisphere has tended to be the focus of study and is known to account for as much as 50% of variance in prognosis for language outcomes (Busby et al., 2023; Wilson et al., 2023). Historically, approaches to understanding and predicting recovery in PSA involve corroborating characteristics of the injury to the brain (Hartwigsen et al., 2013), the presentation of speech-language impairments (Gilmore et al., 2018), along with brain-based biomarkers of stroke recovery (Bartolomeo & Thiebaut de Schotten, 2016). However, how treatment efforts lead to brain changes that result in recovery of language remains unknown. It is proposed that a model of learning, for both behavioral and neural outcomes, is needed to strengthen approaches to rehabilitation. Considering an estimated 50% of recovery variance is not specified by lesion characteristics, a learning model with behavioral and neural components could provide paramount insight into language rehabilitation. That is, some of the recovery variance may be attributable to connectivity amongst classical language brain regions along with other regions involved in learning and be central to the ability to improve communication.

A foundation for the study of learning in PSA is captured in the long-standing proposition that rehabilitation should center on modifying a behavior through experience-based practice. That practice has roots in behavioral psychology and motor learning theory (Schmidt & Lee, 2005) that Baddeley (1993) formalized for cognitive rehabilitation by arguing that a theory of rehabilitation without a model of learning is a vehicle without an engine. Behavioral interventions for linguistic impairments require relearning as they train restorative or compensatory skills. The principles of experience-dependent neural plasticity are increasingly well-characterized: repetitive practice of newly trained skills drives synaptic strengthening, cortical reorganization, and network-level adaptation in both animal models and in humans (Kleim & Jones, 2008; Raymer et al., 2008). However, relatively little is known about how the specific type of learning engaged (i.e., non-linguistic or linguistic, explicit or implicit, statistical or rule/contingency-based) interacts with the integrity of the brain networks that support it in individuals with post-stroke aphasia. It is only recently that the recognition of learning type as a fundamental component of the aphasia rehabilitation process has been considered (Copland et al., 2021; Dignam et al., 2016; Fridriksson et al., 2022; Menahemi-Falkov et al., 2021; Middleton et al., 2016). To date, prior work has reviewed evidence on learning in PSA (Copland et al., 2021; Fama et al., 2022; Middleton et al., 2016; Peñaloza et al., 2022; Schuchard & Thompson, 2014, 2017; Shaqiri et al., 2018a; Vallila-Rohter, 2017; Vallila-Rohter & Kiran, 2013, 2015), the need to recognize the mechanisms of action for aphasia interventions (Fridriksson et al., 2022), and the influence of learning (or not) on intervention outcomes (Dignam et al., 2016; Middleton et al., 2016). Thus, a model of relearning is needed to inform how rehabilitation is approached to optimize interventions so that experience-based changes in associated brain connectivity result in positive outcomes.

The present study applies models of statistical learning, or learning of complex, hierarchical concepts occurs through implicit-statistical routes (Batterink et al., 2015; Reber, 1967; Seger, 1994). Implicit statistical learning (ISL) is a subset of learning that governs humans’ ability to extract environmental regularities through automatic processes that operate outside of conscious awareness (Conway & Christiansen, 2009). ISL is particularly interesting for study of individuals with aphasia given the lower cognitive demand for use of information that is implicitly learned. Further, there is evidence that individuals with PSA have relatively intact implicit learning ability, given the likelihood of spared brain tissue in the neural circuitry engaged for this type of learning (Ramage et al., 2024; Tagarelli et al., 2019).

A focus on ISL in PSA is not new, and there is support for the notion that individuals with aphasia can learn new words and grammar in statistical learning paradigms (c.f., Cope et al., 2017; Schuchard et al., 2017; Vallila-Rohter & Kiran, 2013). Here ‘grammar’ refers broadly to any rule-governed system, linguistic or non-linguistic, rather than specifically to morphosyntax, consistent with the statistical learning literature’s use of artificial grammar paradigms. For example, individuals with left hemisphere stroke can implicitly extract patterns in serial reaction time tasks (Schuchard et al., 2017), learn patterns in letter strings (Schuchard & Thompson, 2014), segment tri-syllabic words based on statistical probability in synthetic languages (Peñaloza et al., 2015; Shaqiri et al., 2018b), learn non-word to picture associations in cross-situational word learning paradigms (Peñaloza et al., 2017), learn paired-associates for nonverbal characters (Vallila-Rohter & Kiran, 2013), and learn artificial grammars (Cope et al., 2017; Schuchard & Thompson, 2017) at accuracy levels near that of healthy controls. ISL of patterns that follow statistical rules (e.g., transitional probability for word or grammar learning) requires a learner to delineate salient cues in incoming stimuli and to pair that salient input with a relevant response. Learning in these paradigms is reinforced through context-specific practice over repeated learning trials. This process of learning engages neural pathways that change from early to late phases of learning (Kleim & Jones, 2008). For example, one model poses three learning interfaces that are involved in the extraction of salient perceptual features, attaching meaning to those features through integration of previous knowledge with new information, and storing that information for long-term retention (Rodríguez-Fornells et al., 2009). Each of these interfaces relies on different networks of brain regions that are thought to be responsible for acquiring and mastering learned patterns, and these networks are likely central to how individuals respond to speech and language interventions. Furthermore, given that the goal of intervention is generalization of treatment-related language gains, this process may be supported by the automaticity that characterizes implicitly learned patterns (Kovacs et al., 2021; Kovacs & Ashby, 2022). If ISL is relevant to the ability to learn and generalize learned skills, then maximizing the types of stimuli and targeting the brain systems associated with the stages of ISL could change the ways that this type of learning is utilized in aphasia rehabilitation. In fact, this is epistemically similar to the Language Stimulation Approach (Schuell et al., 1964). Schuell’s clinical intuition that exposure without explicit teaching supports language recovery anticipated by the ISL principle, though her account invoked a generalized-arousal mechanism, predicting diffuse, non-specific benefit rather than the stage-specific neural systems that ISL frameworks propose. However, as noted previously, the way that the brain relearns and whether the brain networks associated with ISL are spared in PSA is not known.

In an effort to define the brain networks involved in ISL, Ramage and colleagues (2024) used meta-analytic tools to analyze brain imaging data from healthy adults and define an *a priori* set of brain networks engaged in language-based ISL tasks. Meta-analysis with activation likelihood estimation yielded the brain regions most commonly active in functional magnetic resonance (fMRI) studies of non-native and artificial grammar tasks. Meta-analytic connectivity modeling of the connections between those regions, and subsequent hierarchical clustering, identified three subnetworks. These subnetworks were further refined through meta-analytic functional profiling and decoding that indicated their associations with different mental processes across spatial and functional domains. This led to labeling these three subnetworks as relating to language, salience, and cognitive control.

The language (LANG) subnetwork consists of ten brain regions (nodes) including the bilateral inferior frontal pars opercularis (Brodmann area 44), left insula, and two discrete areas in the left inferior frontal triangularis (Brodmann area 45). This subnetwork was found to be commonly active during tasks associated with language processing, including phonological, syntactic, and semantic tasks, particularly with varying cognitive demands. Ramage and colleagues (2024) hypothesized that the LANG network has a central role in language processing but also is conjointly involved with other networks when there is a need to maintain and manipulate incoming language-like stimuli. The LANG subnetwork resembles networks proposed to support ISL, sharing a comparable functional profile by modulating modality-specific inputs during early learning and maintaining that learning within domain-general, higher-level cognition (Frost et al., 2015; Rodríguez-Fornells et al., 2009).

The salience (SAL) subnetwork is made up of 40 nodes including the bilateral insula, frontal, parietal, and cingulate cortex, as well as bilateral caudate nucleus and cerebellar regions. This network is similar to those previously identified as salience or cingulo-opercular networks (Coste & Kleinschmidt, 2016; Seeley et al., 2007) that are thought to be involved in recognition of the unique features of incoming stimuli for conflict resolution and determining correctness. This subnetwork closely aligns with the proposed need in ISL to track regularities and integrate and test potential rules while learning.

The cognitive control (COGCON) subnetwork is a set of 34 nodes spanning the fronto-parietal cortex including bilateral occipital, angular and supramarginal, and middle-superior frontal regions. This network has overlap with the SAL as well as with existing fronto-parietal networks (Cole et al., 2013; Cole & Schneider, 2007) that are often engaged for executive functioning and is associated with tasks involving linguistic and nonlinguistic working memory. It is considered a domain general network engaged as a scaffold for maintenance and application of learned rules.

These meta-analytically derived networks serve here as *a priori* networks of interest to be evaluated for their integrity and relationships to PSA symptomatology using graph theory metrics (Rubinov & Sporns, 2010). Graph theory models brain connectivity as a graph, with regions represented by nodes (e.g., the brain regions of interest) that are connected by edges, or the weighted connections between those nodes. Nodes represent specified volumes within a brain region, and edges are the correlated blood-oxygen-level-dependent resting-state activity between two nodes. In this way, brain networks are characterized by network structure and topology. Graph theory metrics applied to resting-state fMRI data from individuals with PSA have shown that certain networks are more segregated, indicating a break down in connectivity between more distant nodes, and that the amount of segregation relates to accuracy of language performance (Duncan & Small, 2018). Specificity of these networks has also been suggested. For example, cognitive network integrity correlates with verbal executive functioning assessed outside of the scanner, and processing of phonological information with integrity of the language network (Baldassarre et al., 2019). These findings are important in validating the use of resting-state fMRI and graph theory analyses to evaluate brain network integrity and how it relates to relevant behavior, which allows for testing of brain function without having participants perform a task in the scanner.

In this study, differences in connectivity strength or community structure in the ISL *a priori* brain networks (Ramage et al., 2024) are evaluated in resting-state fMRI data from individuals with PSA and compared to that of healthy age-matched control participants. The LANG subnetwork is hypothesized to be most influenced by the presence of a left hemisphere lesion (more of the LANG nodes will be lesioned) and to correspond to aphasia severity (i.e., weaker connectivity, corrected for lesion volume, will correlate with more severe aphasia). Resting-state functional connectivity of networks spatially similar to the SAL and COGCON subnetworks has been found to be predictive of treatment outcomes for individuals with PSA (Falconer et al., 2024). In fact, stronger cognitive control, or domain general, brain network integrity is considered a positive prognostic indicator of outcomes for individuals with aphasia (Fedorenko et al., 2013). Therefore, the hypothesis is that connectivity of the SAL and COGCON subnetworks corrected for lesion volume may be weaker in PSA relative to healthy controls, but their connectivity will be positively correlated with aphasia severity in the chronic stage (i.e., stronger network connectivity will correlate with milder aphasia). These findings will provide insight into the integrity of ISL-related brain networks in individuals with PSA. Because these networks are hypothesized to be important for the extraction and integration of hierarchically organized stimuli, their integrity in PSA and relevance to the language impairment could serve as specific biomarkers related to the language impairment and the potential for the relearning of language in rehabilitation.

## Method

### Participants

Imaging and behavioral data for 30 individuals with aphasia and 18 age-matched healthy controls were reanalyzed from an existing data set described previously (New et al., 2015; Ramage et al., 2020). All participants were between the ages of 18 and 75, were right-handed, and were proficient in English. The participants were without any uncorrected sensory impairment and possessed no cognitive deficits or history of substance abuse. Individuals were also able to complete magnetic resonance imaging (MRI) data acquisition. Healthy control participants performed within normal limits on the Mini-Mental Status Examination (Folstein et al., 1975). The study procedures were conducted in accordance with the Declaration of Helsinki (World Medical Association, 2013) and received ethics approval from the Sydney Local Health Direct Human Research Ethics Committee at Concord Repatriation General Hospital, Sydney, in conjunction with the University of Sydney, Australia. A speech-language pathologist reviewed the participant information statement and consent form with each participant and their care partner, using comprehension questions to confirm that study details were understood before written consent was obtained. Written informed consent was collected from every participant. Demographic information is reported in Table 1.

**Table 1.** Means ± standard deviations for variables characterizing the sample. The groups did not differ for age or years of education, but there were more females in the control than the PSA group (*X*^2^1 = 9.262, *p* = 0.002).

|  | **PSA** | **Control** |
|---|---|---|
| n | 30 | 18 |
| Age in Years | 63 ± 10 | 63 ± 9 |
| Sex (female) | 4 | 10 |
| Education in Years | 15 ± 3 | 16 ± 4 |
| Months post Onset of Stroke | 30 ± 26 |  |

### Language Measures

PSA participants completed a battery of language and speech evaluations including the *Western Aphasia Battery-Revised* (WAB-R) (Kertesz, 2007). For the present study, only the WAB-R scores are reported, but other measures were analyzed and reported in New et al. (2013) and Ramage et al. (2020). Scores are reported in Table 2.

**Table 2.**
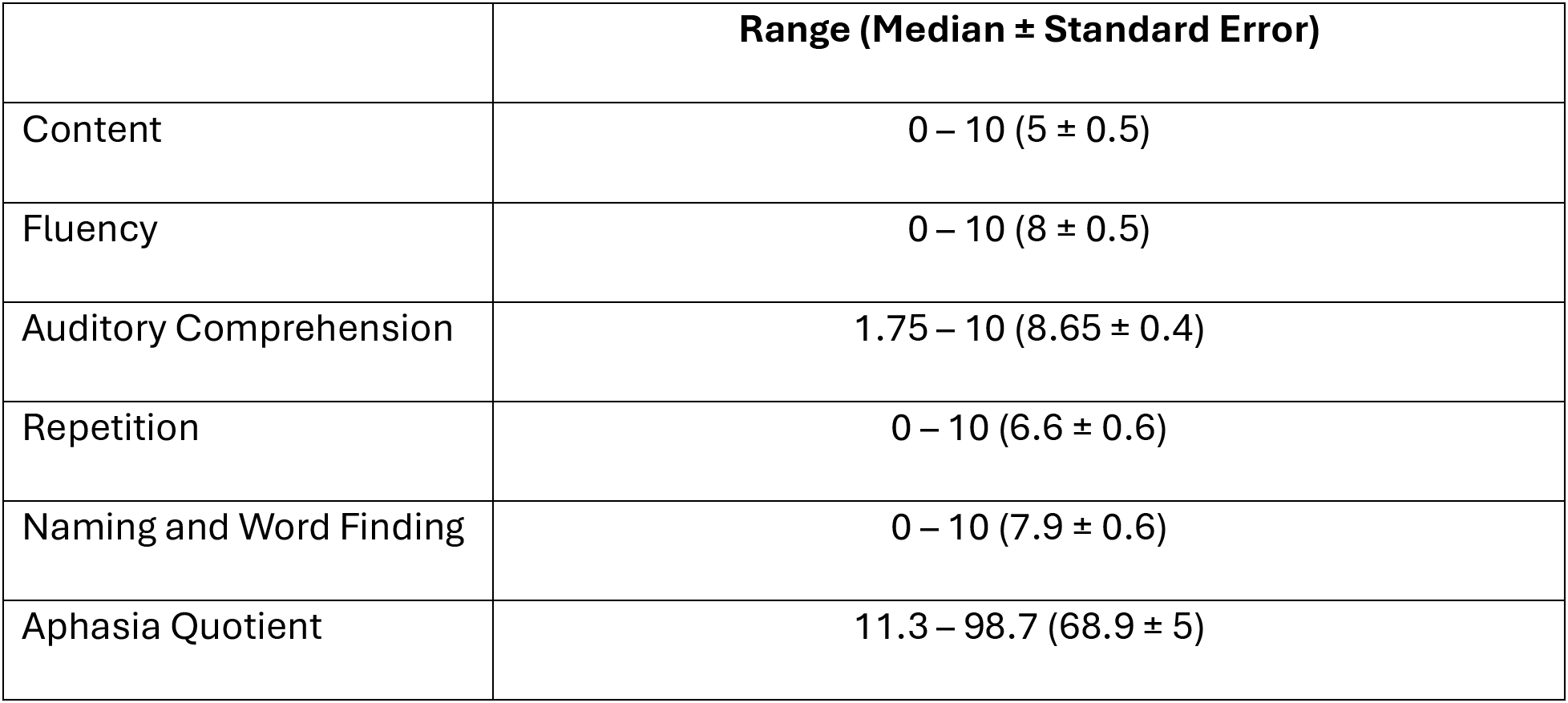
Western Aphasia Battery-Revised (WAB) scores for the PSA group demonstrate a wide range of aphasia severity and difficulty across all subtests. The 30 PSA participants’ aphasia classifications included 11 characterized as nonfluent (7 Broca, 1 Transcortical Motor, 3 Global) and 19 as fluent (9 Anomic, 1 Transcortical sensory, 5 Conduction, 1 Wernicke, and 3 latent). Comorbid apraxia of speech or dysarthria was present for 15 of the participants.

|  | **Range (Median ± Standard Error)** |
|---|---|
| Content | 0 – 10 (5 ± 0.5) |
| Fluency | 0 – 10 (8 ± 0.5) |
| Auditory Comprehension | 1.75 – 10 (8.65 ± 0.4) |
| Repetition | 0 – 10 (6.6 ± 0.6) |
| Naming and Word Finding | 0 – 10 (7.9 ± 0.6) |
| Aphasia Quotient | 11.3 – 98.7 (68.9 ± 5) |

### MRI data acquisition and Imaging Preprocessing

T1-weighted structural and resting-state echo planar fMRI images were collected using a Philips 3T TX MEI Scanner. BOLD contrast was employed to acquire 216 images (TR = 2.2ms, TE = 30ms, flip angle = 90°, in-plane resolution = 3.1 x 3.1 mm^3^, 36 axial slices covering the whole brain). Resting-state echo planar MR images were corrected for slice timing and head motion and normalized spatially to standard Montreal Neurological Institute (MNI) space utilizing SPM8 (Ashburner & Friston, 2005). Any physiological noise was removed to prevent inaccurate correlations. A tissue class for lesions was added to the segmentation to ensure separation of lesions from brain tissue and cerebrospinal fluid. Lesion volumes for all PSA participants were determined using automated lesion identification algorithm (ALI toolbox) in SPM8 (Seghier et al., 2008). This method has been used repeatedly in analyses of stroke data and has been shown to be accurate (Seghier et al., 2016). Lesion volume (in mm^3^) was calculated for each participant and was included as a covariate in all analyses. The extent of the lesions in the sample included frontal, temporal, parietal, and subcortical regions with peak overlap of lesions (23 participants) in the left posterior insula and body of the caudate nucleus (see (Ramage et al., 2020) for visualization of a lesion overlap map for these participants).

### Region of Interest (ROI) Selection

ROI’s were selected based on a previous meta-analysis of 25 fMRI studies of artificial and non-native grammar learning tasks in healthy adults reported in Ramage et al. (2024). Distinct 5mm spherical ROIs for each of the nodes of these networks were created, generating 84 nodes (10 LANG, 40 SAL, and 34 COGCON). The spatial distribution of the nodes is presented in Figure 1, and their coordinates and location labels are provided in Supplemental Materials.

**Figure 1.**
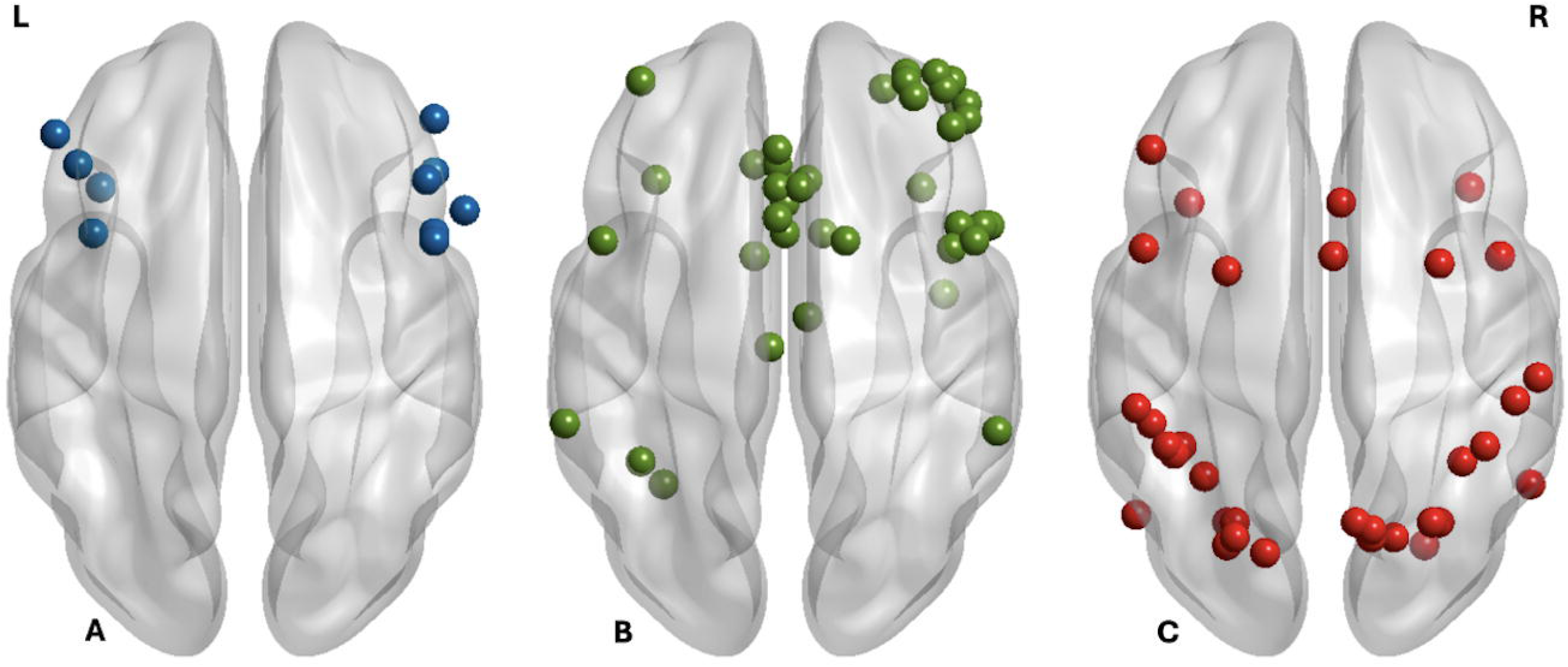
Nodes constituting the *a priori* implicit-statistical learning networks -language (A blue), salience (B green), and cognitive control (C red) subnetworks identified through coordinate-based meta-analytic and functional decoding processes reported in Ramage et al. (2024). Graphic generated in BrainNet Viewer (Xia et al., 2013).

### Data Analysis

#### Resting State Functional Connectivity

Time courses for each of the nodes were extracted for all participants using REX (Duff et al., 2007). Node-by-node correlation matrices for ROIs within each network were computed in MATLAB (The Mathworks Inc., 2016) for each participant and concatenated across participants representing within-network connectivity for each group.

#### Network Structure

##### Within-Network Connectivity

Network-Based Statistics (NBS, Zalesky et al., 2010) was used to test hypothesized differences in connectivity strength between the PSA and Control groups. Additionally, NBS was used to characterize the network structure in terms of the presence or absence of significant edges between subnetwork nodes. NBS is a statistical approach that tests the null hypothesis for interconnected edges, rather than at each connection, by identifying suprathreshold edges between nodes. In this way, NBS offers more statistical power than, for example false discovery rate, to identify edges within networks. Because NBS constructs its null distribution via permutation of group labels rather than relying on equal-sample-size assumptions, it maintains valid family-wise error control even when comparing groups of unequal size, as was the case for the PSA and Control groups here (Zalesky et al., 2010); statistical power, however, remains constrained by the size of the smaller group. Main effects for group differences across subnetworks were assessed across a range of *t-*thresholds to ensure reliability of the results with *t* > 4 (10,000 permutations) and *p* < 0.001 to provide robust findings across a uniform threshold. Effect sizes were calculated with Cohen’s *d* as the average of the significant edges / square root of *n* squared. NBS was also used to identify relationships between functional connectivity, indexed by edge strength, for the 3 hypothesized subnetworks and aphasia severity by adding the WAB-R Aphasia Quotient as a variable in the design matrix.

##### Between-Network Coupling

Between-network connectivity was computed by creating a matrix representing all three subnetworks and extracting and correlating their means by subject (e.g., LANG-and-SAL, LANG-and-COGCON, SAL-and-COGCON) in MATLAB (The Mathworks Inc., 2016). Between-network coupling values were then contrasted for group differences in individuals with PSA relative to controls using the Mann-Whitney U Test in SPSS version 29.0.1.0 (IBM Corporation, 2023) with alpha set at *p* < 0.05, Bonferroni corrected for multiple comparisons. Correlations for between network coupling and WAB-R aphasia quotient in the PSA group were also computed, with statistical significance considered at *rho* > |0.30| and *p* < 0.01.

## Results

Results are first presented for each of the subnetworks separately with characterization of the average connectivity strength and network structure, network-based statistics for edge strength, and associations of edge strength with aphasia severity. Then, the between-network coupling results are reported.

### Network Structure - Within-subnetwork Connectivity

#### Language Subnetwork

Average connectivity was stronger in the healthy controls than the PSA participants for the LANG subnetwork (*z* = -3.237, *p* = 0.001, PSA mean = 0.32, s.d. = 0.16; Control mean = 0.53, s.d. = 0.24), but there were no significant group differences for any single network edge (NBS *p*s > 0.05). Both participant groups had above-threshold bilateral connections between frontal and insular nodes, but there were fewer in the PSA group (n = 27) than the controls (n = 41). As well, the node-to-edge ratios were more evenly distributed in the healthy control than PSA group. The edge ratios in individuals with PSA were highest for three nodes of the right middle and inferior frontal gyri and a node in the left insula (Figure 2).

**Figure 2.**
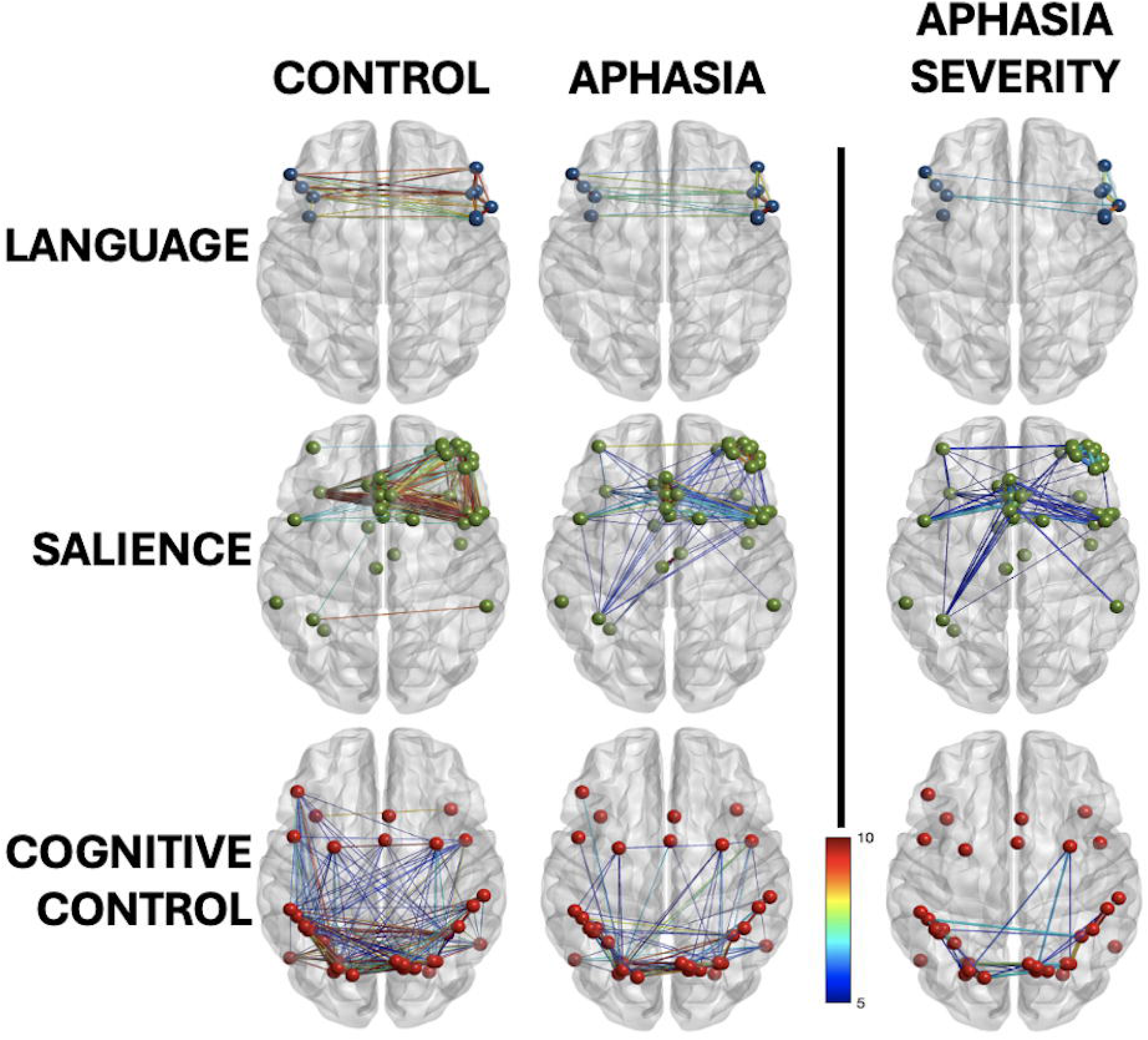
Control (left) and post-stroke aphasia (right) connectivity for the three subnetworks associated with artificial grammar learning – language (top), salience (middle), and cognitive control (bottom) – and the edges significantly correlated aphasia severity (far right) for each. The post-stroke aphasia group had fewer connections than the controls for all three, but average connectivity strength only differed significantly for the language and salience subnetworks. Graphic generated in BrainNet Viewer (Xia et al., 2013).

Severity of aphasia (WAB-R AQ) positively associated with connectivity strength amongst the right middle frontal nodes, specifically those inter-connecting the right middle frontal gyri (*t* = 9.65, *p* < 0.001) and the right inferior frontal operculum and the right middle frontal gyrus (*t* = 6.14, *p* < 0.001). Only two of the inter-nodal connections in the left hemisphere significantly related to AQ – the two nodes within the left inferior frontal triangularis (*t* = 5.15, *p* < 0.001) and the left insula and left inferior frontal triangularis (*t* = 4.39, *p* < 0.001) (Figure 2, top row, right column).

#### Salience Subnetwork

Average connectivity strength within the SAL was stronger in the healthy control than the PSA group (*z* = -3.258, *p* = 0.001, PSA mean = 0.31, s.d. = 0.13; Control mean = 0.52, s.d. = 0.25). This difference was particularly evident upon visual inspection of the network maps, with almost half the number of above-threshold edges in the PSA group (193, Cohen’s *d* = 0.26) than the controls (321, Cohen’s *d* = 0.43) (Figure 1). Distribution of edge ratios was similar across groups, though the left-sided supramarginal and lateral occipital nodes that were well connected in the control group were weaker in the PSA group.

The SAL edges with connection strength related to aphasia severity (WAB AQ) included interconnections between the right frontal pole nodes (*t* = 11.17, *p* < 0.001), right middle frontal and precentral nodes (*t* = 11.03), left superior frontal and paracingulate nodes (*t* = 10.69, *p* < 0.001), and the bilateral caudate nuclei nodes (*t* = 7.5, *p* < 0.001) (Figure 2, middle row, right column).

#### Cognitive Control Network

Average network connectivity strength across the COGCON was weaker in the PSA group relative to the control group (*z* = -2.151, *p* = 0.03, PSA mean = 0.38, s.d. = 0.14; Control mean = 0.49, s.d. = 0.20) but the difference did not survive correction for multiple comparisons. Additionally, no significant group contrast for edge strength was found (all *p*s > 0.05). However, there were fewer above-threshold edges in the PSA group (193 edges, Cohen’s *d* = 1.41) than the Controls (321 edges, Cohen’s *d* = 1.84). The strongest edges in both groups were amongst the parieto-occipital nodes bilaterally, and the reduced number of edges was most evident in the frontal connections on the left. Edge ratios were similar in both groups. The edges with connection strength that related to aphasia severity included the interconnections amongst the left superior parietal lobule and supramarginal gyrus (t > 11, p < 0.001).

The COGCON edges with connection strength associated with aphasia severity (AQ) were those amongst the left superior parietal lobule and supramarginal gyrus. These included 2 connections amongst superior parietal lobule nodes (*t* = 12.24, *p* < 0.001) and (*t* = 11.34, *p* < 0.001), the superior lobule and the supramarginal gyrus (*t* = 11.35, *p* < 0.001), and amongst nodes in the supramarginal gyrus ( *t* = 11.09, *p* < 0.001).

### Network Topography

#### Between Network Results

The groups differed for between-network coupling strength for LANG-and-SAL (*z* = -3.29, *p* = 0.002, PSA mean = 0.32, s.d. = 0.14; Control mean = 0.55, s.d. = 0.27), LANG-and-COG (*z* = -3.13, *p* = 0.001, PSA mean = 0.32, s.d. = 0.16; Control mean = 0.50, s.d. = 0.25), and SAL- and-COG (*z* = -3.10, *p* = 0.002, PSA mean = 0.32, s.d. = 0.15; Control mean = 0.49, s.d. = 0.24). That is, the correspondence of activity between the networks was greater in the healthy control group than the PSA group.

#### Between-Network Coupling Correlation with Aphasia Severity

The average connectivity across all nodes of the LANG network correlated with the WAB AQ (*rho* = -0.38, *p* = 0.04); such that the stronger the connectivity amongst these nodes at rest, the milder the aphasia. No significant correlations existed between aphasia severity and coupling strengths between the three networks (all *k* < |0.30|), suggesting that while coupling was generally weaker in the PSA group, it did not account for variance in aphasia severity.

## Discussion

Relearning is an essential skill to recovery and the neural reorganization that supports positive adaptation following brain damage. The focus of intervention typically begins by limiting the severity of the initial injury, followed by rehabilitation to reorganize the brain to restore and compensate for impaired function. Brain damage to the language network is very likely to disrupt the coordinated activation across multiple networks that experience-dependent learning of linguistic patterns requires, rather than simply impairing language processing in isolation. These alterations may be best elucidated through assessment of neural network configuration and connectivity – both within-network and between-network (network coupling). As established in the Introduction, individuals with PSA show evidence of ISL across a range of paradigms. What remained unknown prior to the present study was whether ISL-associated brain networks identified in healthy adults are preserved in this population and whether their integrity corresponds to individual differences in language impairment.

What remains unknown is how rules or patterns from ISL may generalization to untrained stimuli or contexts in some individuals with PSA. What is known is that, in the few studies investigating generalization of rules learned in artificial grammars, it does not occur, specifically in those with nonfluent aphasia profiles (Cope et al., 2017; Hoen et al., 2003). Note that these studies examined ISL recognition and discrimination performance with varying levels of exposure; and thus, the limitation reflects a variable ISL input-processing capacity. Whether these limitations reflect reduced ISL capacity, lesion-related disruption of the networks supporting it, or other factors related to aphasia severity remains unclear. As such, this study aimed to determine how the brain networks associated with ISL are altered in PSA and whether those alterations are relevant to the language impairment.

Previous work utilized coordinate-based meta-analytic tools to identify three brain networks commonly recruited for ISL of artificial grammars in healthy adults and that were uniquely associated with language, salience, or cognitive control (Ramage et al., 2024). The hypothesis is that the ISL language subnetwork would be disrupted in participants with PSA and that the extent of disruption would correlate with aphasia severity. Further, it is hypothesized that the salience and cognitive control networks may not differ from healthy participants. This would provide some support for the idea that these networks may be optimized to support learning in treatment approaches based in ISL in a way similar to that utilized with success in childhood language disorders (c.f., Alt, 2018; Alt et al., 2012).

By using networks of brain regions found to be involved in ISL and differentiated based on their associations with mental operations, it was found that individuals with PSA have weaker connectivity strength amongst all three networks relative to age-matched healthy controls, though this was only significant for the language and salience networks. Specifically, bilateral connectivity was reduced for the language and salience networks in PSA, particularly amongst frontal nodes (Figure 2). In relation to language impairment, strength of internodal connectivity in subsets of edges for all three networks were significantly associated with aphasia severity (WAB-R AQ) suggesting that each may be involved in language impairment to an extent.

The ISL language subnetwork identified in Ramage et al. (2024) is limited to bilateral frontal regions and does not extend to the temporo-parietal cortex that is expected in the classic language network. As found previously in this cohort of individuals with PSA (New et al., 2015; Ramage et al., 2020), inter-hemispheric connectivity in these frontal regions was weaker in the PSA group relative to the controls, and there was stronger internodal connectivity amongst the right frontal nodes than any other supra-threshold edges. Further, it was the connectivity strength amongst these right frontal nodes that correlated with the WAB-R AQ such that those with stronger connectivity had less severe language impairment. Previous reports of right hemisphere recruitment following left hemisphere stroke have suggested that such a shift in language dominance is maladaptive when seen in task-evoked fMRI (Klingbeil et al., 2019; Saur et al., 2006; Wilson & Schneck, 2021). However, these findings used task-evoked neural activity, which reflects the attempt to use the right hemisphere to *produce* language when the left hemisphere is damaged. When the right hemisphere attempts language production, performance is typically limited (c.f., Benson & Zaidel, 1985), but right hemisphere contributions to language comprehension and pattern recognition are less well characterized and may be more substantial. Resting-state connectivity of the right frontal cortex in left hemisphere stroke has only been noted as hypo-connected relative to healthy controls (Sandberg, 2017), and not related to other individual factors like aphasia severity. Some resting state and task-evoked fMRI studies of speech auditory feedback in individuals with stroke have shown changes in right hemisphere functional and effective connectivity reflecting cortical reorganization (Behroozmand et al., 2015; Sarmukadam et al., 2025; Teki et al., 2013). Further, in this sample of PSA participants, stronger inter-hemispheric connectivity between frontal regions also correlated with severity of apraxia of speech (New et al., 2015), and thus the findings are likely not specific to language processing, but nonetheless have relevance to communication impairment and recovery post stroke. Similarly, Yourganov et al. (2021) report increased resting-state connectivity in right hemisphere regions associated with speech and sensorimotor functions. Given that the WAB-R AQ includes production-heavy subtests that are directly affected by comorbid apraxia of speech, and that 15 of the 30 PSA participants had comorbid apraxia of speech or dysarthria, the right frontal connectivity-AQ correlation must be cautiously interpreted. Replication in a cohort without comorbid motor speech disorders, or using aphasia measures less sensitive to production demands, would help clarify whether this connectivity pattern is specific to language processing, motor speech, or domain-general communication. It is also notable that ISL, as operationalized in the present study and the meta-analyses from which the networks were derived, is fundamentally an input-processing capacity – recognition of statistical regularities in incoming stimuli. Whether the right frontal network integrity that correlates with milder aphasia severity also supports the deployment of implicitly learned patterns in spoken output is an open question to be addressed in future work.

Salience network connectivity also differed between the groups, for average network connectivity strength, for the numbers of super-threshold edges, and for the spatial distribution of those edges. Visual inspection of the connectivity maps (Figure 2) demonstrates the most obvious differences around specific nodes (right paracingulate and frontal and left superior frontal) having more connections in the controls. Salience subnetwork edges that associated with aphasia severity involved bilateral nodes in paracingulate cortex and the bilateral caudate nuclei nodes (Figure 2, middle row, rightmost column). These medial brain structures are part of an integrated fronto-subcortical circuit that is thought to support higher-order language necessary for conflict resolution and suppression of linguistic alternatives, found in healthy controls who are mono- or bilingual (Copland et al., 2021; Li et al., 2016; Tomasi & Volkow, 2012). In individuals with PSA, these regions are thought to mediate or moderate domain-general cognitive control and compensatory neuroplasticity necessary for monitoring and suppressing errors (Anwia et al., 2026; Falconer et al., 2024).

Connectivity of the cognitive control network did not differ between groups, though the spatial structure of the network appeared somewhat different between groups upon visual inspection. Markedly, in the PSA group there was less integration of connections amongst nodes in the frontal and parietal cortex, bilaterally (Figure 2, bottom row). This finding would support those of Federenko and colleagues suggesting that this network, akin to their multiple demand network, does not play a role in the language impairment (particularly for auditory comprhension) in aphasia (Billot et al., 2026; Fedorenko, 2014; Fedorenko et al., 2013). However, some edges of the cognitive control subnetwork were associated with aphasia severity (specifically the interconnections between and within the left and right superior parietal lobules).

To address potential inter-reliance between the subnetworks, coupling between the three ISL subnetworks was assessed. Strength of the coupling differed significantly between groups with a higher level of integration (stronger coupling) in the control participants. However, coupling strength did not correlate with aphasia severity. This is similar to that found by Billot et al. (2026), who reported little coupling between their multiple demand and language networks and that integration between the networks was not relevant to the language profiles of their PSA participants. Other studies investigating brain network coupling findings relevant to language processing or aphasia severity assessed brain connectivity during task-based fMRI (Geranmayeh et al., 2014, 2017), suggesting that this aspect of network correlation differs in the resting-state and task-evoked condition, thus warranting further study.

## Limitations

This study represents a reasonable first step toward linking domain-general learning mechanisms to post-stroke language, but there are limitations that should inform interpetation of the findings. First, a network-based reverse inference approach was used in which meta-analytically defined ISL networks served as *a priori* regions of interest to test network integrity and severity associations in PSA. The *a priori* networks were derived from coordinate-based meta-analyses of artificial and non-native grammar learning studies, i.e., ISL was not measured in the stroke sample. The PSA cohort was one of convenience, recruited from an existing clinical dataset rather than assembled specifically to test ISL; as such, direct behavioral measurement was not feasible within the scope of the study. And given the difficulties of assessing ISL during scanning, or language-like processing in this population given potential confounds of cognitive and language impairments, resting-state network integrity offeres a practical and theoretically-motivated entry point to test this relationship in this population. Future work with prospective designs and behavioral implicit learning tasks adapted for use in aphasia would help to confirm and extend the current findings.

As well, the use of resting-state functional connectivity as a proxy for network integrity introduces its own inferential distance from the task-based activation patterns used to define the ISL subnetworks. Resting-state and task-evoked activation are related but not interchangeable. Resting-state approaches have the advantage of being tolerable and feasible in stroke patients with limited comprehension and endurance, and prior work has shown that resting-state network organization meaningfully predicts task-related function across a range of domains (Smith et al., 2009). The findings reported here should be understood as evidence of a structural/functional substrate consistent with the proposed mechanism (i.e., ISL), not a direct confirmation.

Aphasia severity was indexed by WAB-R AQ, but the construct is multiply-determined, being influenced by lesion size and location, white matter tract disruption, premorbid cognitive and linguistic ability, time post-onset, and many other factors. Without a measure of ISL in these participants, it is possible that associations between network integrity and aphasia severity are driven by shared variance with other lesion-related or demographic factors, rather than a mechanistic link to implicit learning capacity. Without a behavioral measure of ISL in these participants, it is not possible to isolate it from shared variance with these other factors. Nonetheless, the robust association between a theoretically specific subnetwork and clinical severity, even without fully partialling out all potential confounds, is a meaningfu step that can build more hypothesis-driven work (e.g., incorporating lesion-symptom mapping alongside behavioral implicit learning measures (e.g., (Lim et al., 2019)) to estabish specificity.

Finally, the meta-analytic connectivity maps were derived from neurologically healthy populations and their anatomical and functional correspondence in a lesioned, reorganizing brain is yet nascent. Nonetheless, applying networks defined independently of the current sample avoids circularity in network selection, and provides a hypothesis-driven basis for testing network-severity relationships, rather than relying on data-driven networks that risk overfitting to the current dataset.

Despite these limitations, this study provides preliminary evidence that ISL networks are relevant to post-stroke language outcomes, offering a novel, mechanistically motivated framework for understanding individual variability in aphasia severity and a foundation for future studies directly testing ISL in this population.

## Conclusions

The study hypotheses were that weaker language subnetwork connecitvity would correspond with more severe aphasia, and that salience and cognitive control subnetwork connectivity would not. The findings were more nuanced, suggesting specificity by laterality in the language subnetwork – stronger connections between right frontal regions corresponded to milder aphasia. In fact, for all subnetworks, stronger connectivity amongst right frontal nodes associated with milder aphasia. The language subnetwork findings may be consistent with the theory put forth by Saur and colleagues (2006) that laterality across recovery occurs in three phases: (1) reduced activation of the left-hemispheric language network following stroke, (2) upregulation of parallel network in the right hemisphere, followed by (3) a decrease in right-sided activation and a shift back to the left-sided dominance for language (Saur et al., 2006). Failure to recover left hemisphere language dominance has typically been considered maladaptive, given that the right hemisphere is not specialized for language processing. However, that right frontal connecitivity is associated with better aphasia outcomes supports the importance of mental processes outside of the classical language cortex, particularly in the right middle and inferior frontal cortex, as has been found previously (Heiss & Thiel, 2006; Kiran et al., 2019; Turkeltaub et al., 2012). These findings imply that right frontal connectivity may be associated with compensatory rather than maladaptive responses to injury. Importantly, though, is that bilateral caudate connectivity also associated with aphasia severity, highlighting that interconnections amongst regions outside of the traditional language areas and their right hemisphere homologs are contributing to individual variance in the impairment. As such, this study contributes to understanding about the right frontal and subcortical mental processes that contribute to recovery, including not only right-sided homologs of language areas, but also those that underly ISL. Further study, including measurement of both ISL and functional connectivity in individuals with PSA will elucidate the relevance of the elements of learning that may be beneficial.

## Acknowledgments

The authors acknowledge the contributions of members of the UNH Cognition, Brain, and Language Team for their help in organization and analyses of the data. They also thank Cathy Price and the Predicting Language Outcomes and Recovery after Stroke laboratory for conducting the automated lesion identification for this project.

