## Supplemental Materials for "Connectivity of Implicit Statistical Learning Networks is Associated with Language Impairment Severity in Post-Stroke Aphasia"

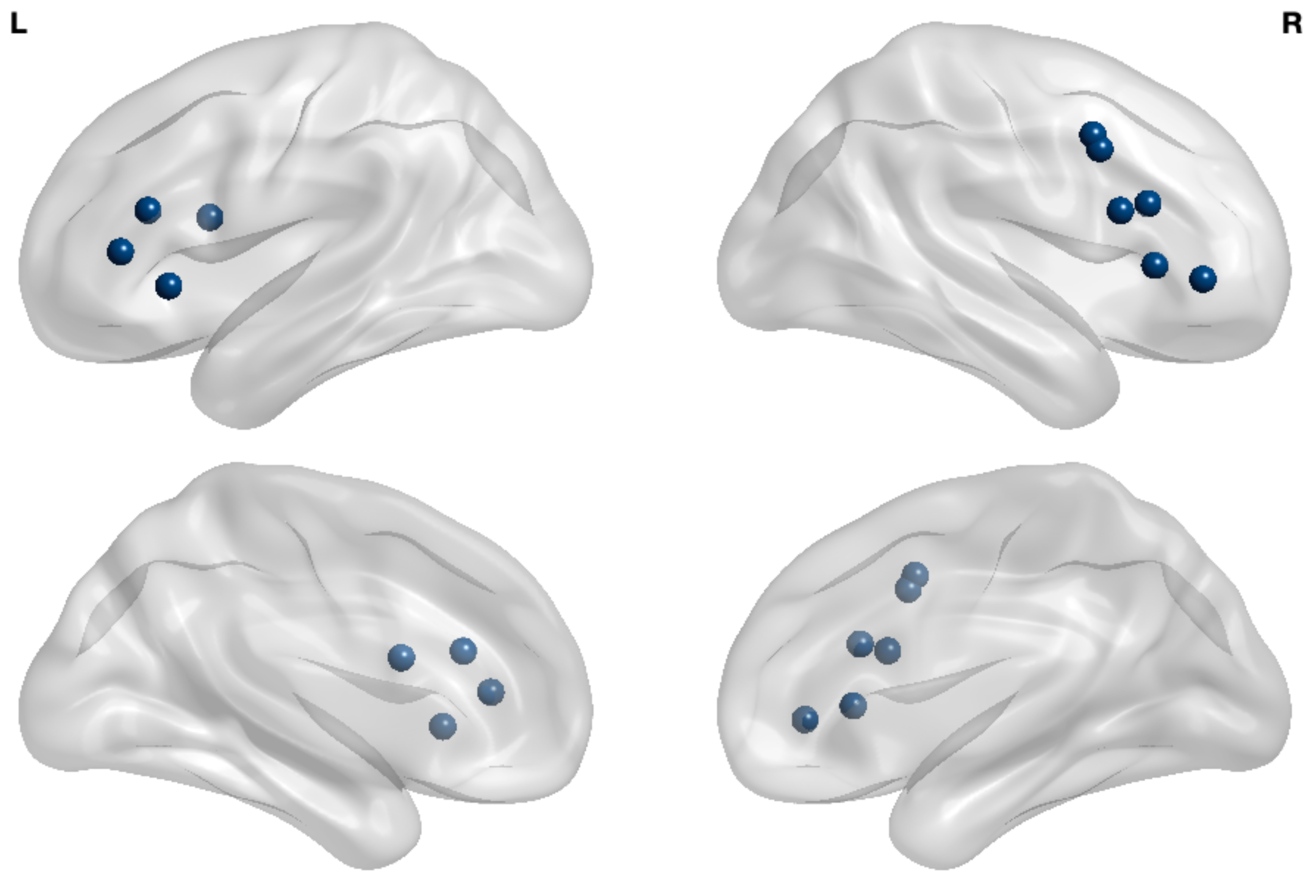
**Supplemental Figure 1**. The ISL Language subnetwork is made up of 10 nodes including the middle and inferior frontal gyri bilaterally.

| Node | Label | x | y | z | Macroanatomy | Cytoarchitecture |
| --- | --- | --- | --- | --- | --- | --- |
| 1 | rIFGtri1 | 50 | 26 | 4 | 34% IFG pars triangularis; 6% IFG opercularis | 45/44/OP8/OP9 |
| 2 | rMFG2 | 48 | 24 | 22 | 20% MFG, 18% IFG pars opercularis | 45/44 |
| 3 | rIFGop3 | 58 | 16 | 20 | 62% IFG pars opercularis; 10% Precentral Gyrus | 44/45 |
| 4 | rFpole4 | 50 | 40 | 0 | 70% Frontal Pole; 12% IFG triangularis | 45 |
| 5 | rMFG5 | 50 | 8 | 42 | 41% MFG, 30% Precentral Gyrus | 44 |
| 6 | rMFG6 | 50 | 10 | 38 | 40% MFG, 28% Precentral | 44 |
| 7 | lIFGop7 | -40 | 10 | 18 | 41% IFG opercularis, 10% MFG | 44 |
| 8 | lIns8 | -38 | 22 | -2 | 36% Insula, 25% Frontal Orbital Cortex | Id7 |
| 9 | lIFGtri9 | -50 | 36 | 8 | 43% IFG triangularis; 31% Frontal Pole | 45 |
| 10 | lIFGtri10 | -44 | 28 | 20 | 44% IFG triangularis; 25% MFG | 45 |

**Supplemental Table 1**. Language subnetwork nodes and coordinates as well as the macroanatomy and cytoarchitecture (Zilles et al., 2002; Zilles & Amunts, 2010) descriptions from the SPM Anatomy Toolbox 3.0 (Eickhoff et al., 2005, 2006, 2007).

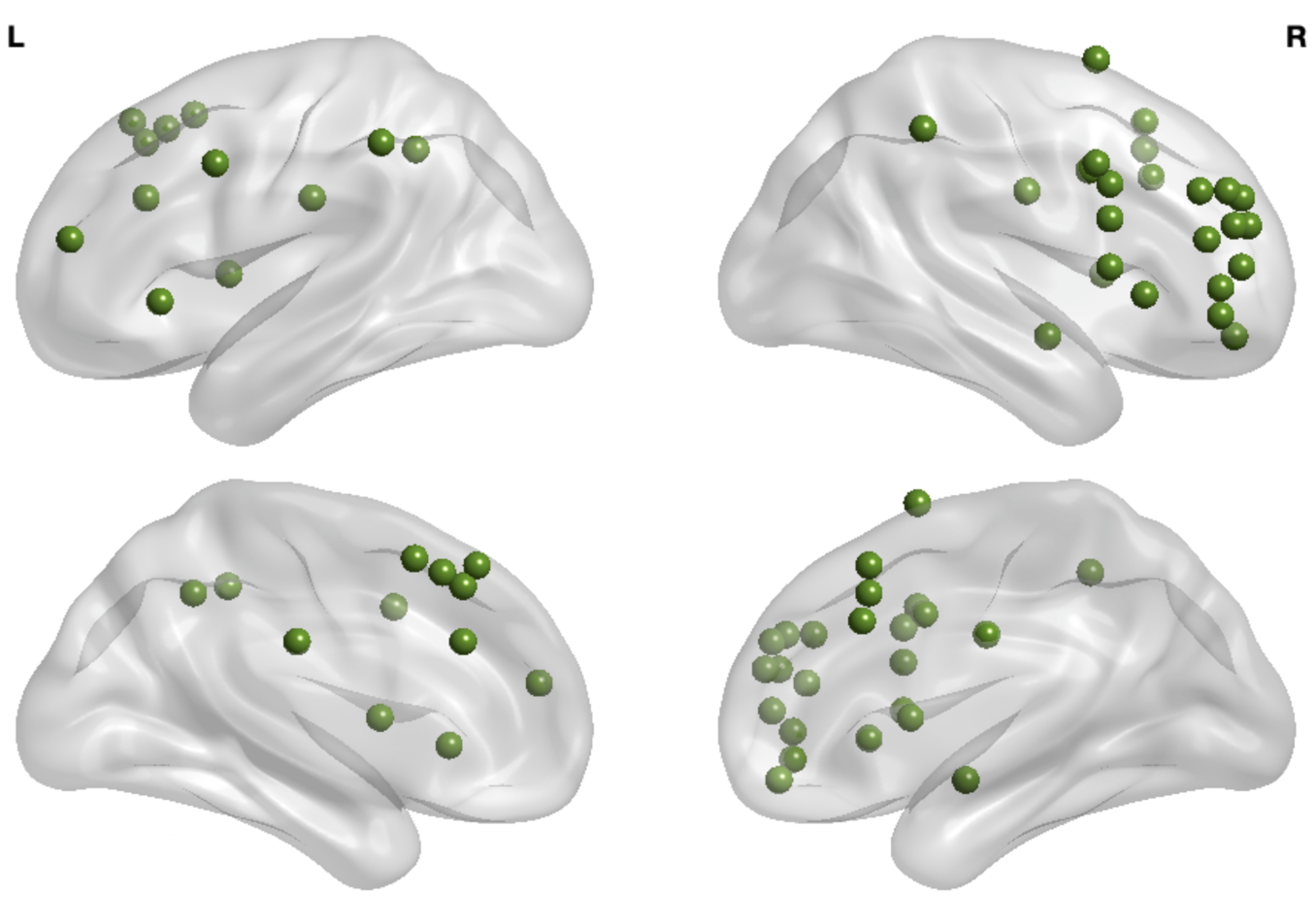

**Supplemental Figure 2**. The ISL Salience subnetwork is made up of 40 nodes including the frontal, parietal, temporal, and subcortical structures bilaterally.

| Node | Label | x | y | z | Macroanatomy | Cytoarchitecture |
| --- | --- | --- | --- | --- | --- | --- |
| 1 | rParacing1 | 6 | 24 | 34 | 50% Paracingulate Gyrus; 36% Cingulate Gyrus (anterior division) |  |
| 2 | rSFG2 | 4 | 22 | 50 | 60% SFG, 26% paracingulate gyrus |  |
| 3 | lSFG3 | -2 | 22 | 48 | 35% SFG, 30% paracingulate gyrus |  |
| 4 | Paracing4 | 0 | 18 | 46 | 51% Paracingulate Gyrus, 4% SFG |  |
| 5 | rParacing5 | 2 | 22 | 42 | 71% Paracingulate Gyrus, 9% Cingulate (anterior) |  |
| 6 | lParacing6 | -2 | 14 | 52 | 36% Paracingulate Gyrus, 24% SFG | 6mr/preSMA |
| 7 | lSFG7 | -2 | 28 | 44 | 34% SFG, 31% Parcing |  |
| 8 | lParacing8 | -8 | 28 | 28 | 44% Paracingulate Gyrus, 39% Cing (anterior) |  |
| 9 | Cing9 | 0 | 10 | 28 | 87% Cingulate Gyrus (anterior) |  |
| 10 | lSFG10 | -2 | 32 | 50 | 73% SFG |  |
| 11 | rINS11 | 36 | 22 | 0 | 47% Insula, 16% Frontal Operculum | id7, 45 |
| 12 | rIFG12 | 54 | 12 | 8 | 58% IFG; 18% Precentral Gyrus | 44 |
| 13 | rIFGop13 | 52 | 12 | 22 | 48% IFG pars Opercularis; 25% Precentral gyrus | 44 |
| 14 | rMFG14 | 44 | 6 | 36 | 33% MFG, 30% Precentral Gyrus | 44 |
| 15 | rPcnt15 | 50 | 8 | 38 | 42% Precentral Gyrus, 34% MFG | 44 |
| 16 | rMFG16 | 46 | 12 | 32 | 28% MFG, 21% Precentral Gyrus | 44 |
| 17 | lINS17 | -34 | 24 | -2 | 38% Insula; 34% Frontal Orbital Cortex | id7 |
| 18 | rFpole18 | 44 | 50 | 8 | 86% Frontal Pole |  |
| 19 | rFpole19 | 44 | 38 | 30 | 61% Frontal Pole; 29% MFG |  |
| 20 | rFpole20 | 40 | 52 | 20 | 74% Frontal Pole |  |
| 21 | rFpole21 | 48 | 44 | 2 | 90% Frontal Pole; 2% IFG p triangularis |  |
| 22 | rFpole22 | 42 | 48 | 20 | 85% Frontal Pole |  |
| 23 | rFpole23 | 32 | 52 | 20 | 83% Frontal Pole |  |
| 24 | rFpole24 | 32 | 50 | 28 | 86% Frontal Pole |  |
| 25 | rFpole25 | 34 | 46 | 30 | 85% Frontal Pole |  |
| 26 | rFpole26 | 48 | 40 | 16 | 71% Frontal Pole; 11% MFG |  |
| 27 | rFpole27 | 46 | 44 | -6 | 91% Frontal Pole |  |
| 28 | rSMG28 | 56 | -42 | 48 | 66% Supramarginal Gyrus; 20% Angular Gyrus | PFm (IPL), PF(IPD); hIP2(IPS) |
| 29 | lSMG29 | -58 | -40 | 44 | 44% Supramarginal Gyrus (44% anterior division, 34% posterior division) | PF (IPL); hIP2(IPS); PFm(IPL) |
| 30 | lFpole30 | -38 | 50 | 16 | 85% Frontal Pole |  |
| 31 | rCaud31 | 10 | 10 | 6 | 91% Right Caudate |  |
| 32 | lSMG32 | -38 | -50 | 42 | 25% Suprmarginal Gyrus (posterior division); 21% Angular Gyrus | hIP1(IPS); hIP2(IPS), hIP3(IPS), 7PC(SPL) |
| 33 | lCaud33 | -8 | 4 | 6 | 53% Left Caudate |  |
| 34 | lMFG34 | -48 | 8 | 38 | 33% Middle Frontal Gyrus; 21% Precentral Gyrus | 44 |
| 35 | lCbllm35 | -32 | -56 | -32 | 78% Cerebellum VI; 20% Left Crus I |  |
| 36 | rFpole36 | 26 | 48 | -12 | 58% Frontal Pole | Fo3 |
| 37 | rPP37 | 42 | -6 | -12 | 61% Planum Polare; 15% Insular Cortex | Id1 |
| 38 | lCing38 | -4 | -20 | 28 | 43% Cingulate Gyrus (posterior division); 11% Cingulate Gyrus (anterior division) |  |
| 39 | rSFG39 | 16 | 8 | 68 | 56% Superior Frontal Gyrus | 6d2 |
| 40 | rCing40 | 6 | -12 | 30 | 23% Cingulate Gyrus (anterior); 11% Cingulate Gyrus (posterior) | 33 |

**Supplemental Table 2**. Salience subnetwork nodes and coordinates as well as the macroanatomy and cytoarchitecture (Zilles et al., 2002; Zilles & Amunts, 2010) descriptions from the SPM Anatomy Toolbox 3.0 (Eickhoff et al., 2005, 2006, 2007).

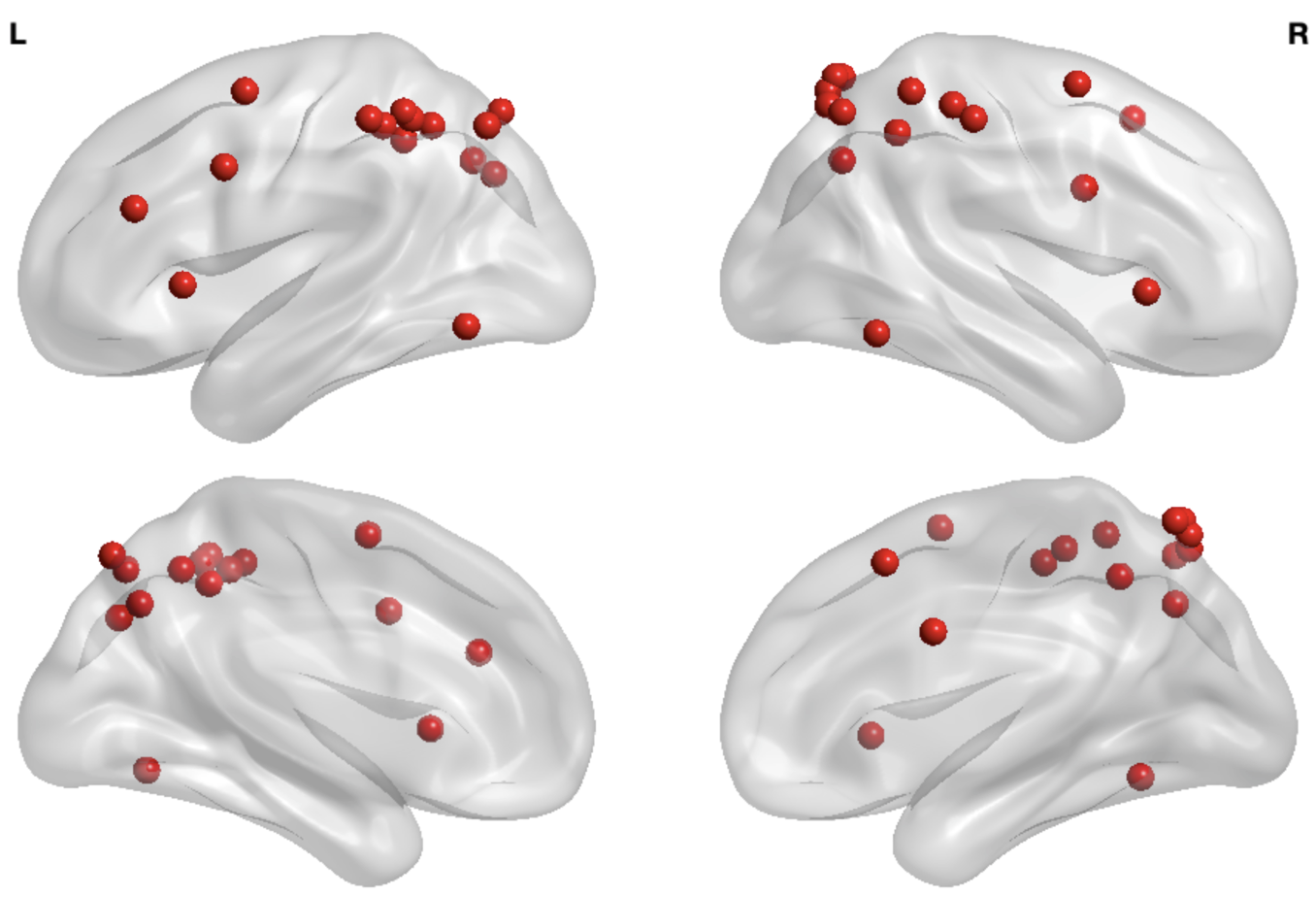

**Supplemental Figure 3**. The ISL Cognitive Control subnetwork is made up of 34 nodes covering the frontal, parietal, and occipital gyri bilaterally.

| Node | Label | x | y | z | Macroanatomy | Cytoarchitecture |
| --- | --- | --- | --- | --- | --- | --- |
| 1 | rLOG1 | 30 | -66 | 38 | 40% lateral occipital cortex; 13% SMG | hIP5, hIP6, hIP8 |
| 2 | rSPL2 | 36 | -50 | 46 | 37% SPL, 26% Angular | hIP3, hIP2, hIP1 |
| 3 | rSMG3 | 50 | -34 | 54 | 30% SMG(ant), 26% SMG(post) | 1, 2, PFt(IPL), PF(IPL), PFm(IPL) |
| 4 | rLOG4 | 28 | -66 | 52 | 60% Lateral Occipital Cortex | 7A, hIP3(IPS) |
| 5 | rSMG5 | 56 | -28 | 50 | 62% SMG(ant), 8% Postcentral | PFt(IPL), PF(IPL), 2, hIP2(IPS), 1 |
| 6 | rANG6 | 42 | -46 | 58 | 57% SPL, 10% Angular | hIP3(IPS), 7PC(SPL), PGa(IPL) |
| 7 | rLOG7 | 18 | -70 | 54 | 52% Lateral Occipital Cortex (sup) | 7P, 7A |
| 8 | rLOG8 | 12 | -70 | 58 | 58% Lateral Occipital Cortex (sup), Precuneus | 7P, 7A |
| 9 | rLOG9 | 12 | -68 | 62 | 59% Lateral Occipital Cortex(sup), Precuneus | 7P, 7A |
| 10 | rLOG10 | 8 | -66 | 62 | 34% Lateral Occipital Cortex, 31% Precuneus | 7P, 7A |
| 11 | lLOG11 | -26 | -72 | 34 | 38% Lateral Occipital cortex; 19% SPL | hIP5, hIP8 |
| 12 | lLOG12 | -24 | -70 | 48 | 58% Lateral Occipital Cortex (sup) | hIP6, 7A, hIP8 |
| 13 | lPcun13 | -24 | -66 | 38 | 54% Lateral Occipital Cortex, Precuneus | hIP8, hIP3, hIP5, hIP6 |
| 14 | lANG14 | -32 | -54 | 48 | 42% SPL, 11% Angular | hIP3, 7PC, hIP1, 7A |
| 15 | lSPL15 | -40 | -48 | 50 | 41% SPL, 22% SMG | hIP3, 7PC, hIP1, hIP2 |
| 16 | lSPL16 | -38 | -46 | 44 | 33% SPL, 32% SMG | hIP1, hIP3, hIP2, 7PC |
| 17 | lSPL17 | -42 | -46 | 52 | 39% SPL, 26% SMG | hIP3, 7PC, hIP1, 5L, hIP2 |
| 18 | lSMG18 | -46 | -40 | 48 | 23% SMG(post); 21% SMG(ant) | 2, PFt(IPL) |
| 19 | lSMG19 | -50 | -36 | 50 | 38% SMG(ant); 18% Postcentral Gyrus | 2, PFt(IPL), 1 |
| 20 | lLOG20 | -16 | -74 | 52 | 70% Lateral Occipital Cortex, 3% Precuneus | hIP8, 7A, 7P |
| 21 | rPcnt21 | 46 | 4 | 30 | 25% Precentral, 24% Frontal Pole | 44 |
| 22 | rITG22 | 54 | -56 | -12 | 71% ITG, 8% MTG (temporooccipital) |  |
| 23 | rLOC23 | -50 | -64 | -10 | 49% Lateral Occipital Cortex; 29% ITG | FG4, FG2 |
| 24 | rMFG24 | 30 | 2 | 60 | 37% MFG, 24% SFG | 6d3 |
| 25 | lMFG25 | -48 | 6 | 36 | 35% MFG; 34% Precentral | 44 |
| 26 | lSFG26 | -26 | 0 | 58 | 33% SFG, 27% MFG | 6d3, 6d2 |
| 27 | lMFG27 | -46 | 32 | 24 | 70% MFG, 12% Frontal Pole | 45 |
| 28 | rOperc28 | 38 | 22 | 0 | 33% Frontal Operculum; 29% Insula | Id7, 45, OP9, OP8 |
| 29 | lCbllm29 | -26 | -72 | -50 | 85% Cerebellum VIIb, 9% Left Crus II |  |
| 30 | lCbllm30 | -26 | -66 | -26 | 96% Cerebellum VI; 3% Crus I |  |
| 31 | rParacing31 | 4 | 18 | 50 | 51% Paracingulate Cortex; 32% SFG | 6mr/pre-SMA |
| 32 | lINS32 | -36 | 18 | 2 | 62% Insula; 16% Frontal Operculum | Id7, OP8, 44 |
| 33 | rCbllm33 | 26 | -72 | -50 | 78% Cerebellum Iib, 15% CrusII |  |
| 34 | rCing34 | 2 | 4 | 30 | 87% Cingulate Cortex (anterior) | 33 |

**Supplemental Table 3**. Cognitive control subnetwork nodes and coordinates as well as the macroanatomy and cytoarchitecture (Zilles et al., 2002; Zilles & Amunts, 2010) descriptions from the SPM Anatomy Toolbox 3.0 (Eickhoff et al., 2005, 2006, 2007).

Eickhoff, S. B., Heim, S., Zilles, K., & Amunts, K. (2006). Testing anatomically specified hypotheses in functional imaging using cytoarchitectonic maps. *NeuroImage*, *32*(2), 570–582. https://doi.org/10.1016/j.neuroimage.2006.04.204

Eickhoff, S. B., Paus, T., Caspers, S., Grosbras, M.-H., Evans, A. C., Zilles, K., & Amunts, K. (2007). Assignment of functional activations to probabilistic cytoarchitectonic areas revisited. *NeuroImage*, *36*(3), 511–521. https://doi.org/10.1016/j.neuroimage.2007.03.060

Eickhoff, S. B., Stephan, K. E., Mohlberg, H., Grefkes, C., Fink, G. R., Amunts, K., & Zilles, K. (2005). A new SPM toolbox for combining probabilistic cytoarchitectonic maps and functional imaging data. *NeuroImage*, *25*(4), 1325–1335. https://doi.org/10.1016/j.neuroimage.2004.12.034

Zilles, K., & Amunts, K. (2010). Centenary of Brodmann’s map—Conception and fate. *Nature Reviews. Neuroscience*, *11*(2), 139–145. https://doi.org/10.1038/nrn2776

Zilles, K., Schleicher, A., Palomero-Gallagher, N., & Amunts, K. (2002). Quantitative analysis of cyto- and receptor architecture of the human brain. In J. Mazziotta & A. W. Toga (Eds.), *Brain Mapping: The Methods* (pp. 573–602). Elsevier.
